# *tidypopgen*: Tidy Population Genetics in R

**DOI:** 10.1101/2025.06.06.658325

**Authors:** Evelyn J. Carter, Eirlys E. Tysall, Jason Hodgson, Andrea Manica

## Abstract

As genome-wide data has become increasingly available, software libraries for their analysis have proliferated. While new tools for downstream analyses are constantly emerging, existing workflows are hindered by inefficiencies. Switching between coding languages and object types in the early stages of pipelines wastes researchers’ time, impedes reproducibility, and creates opportunity for error. To confront these obstacles, we introduce *tidypopgen*, a comprehensive R package for population genetic analysis of biallelic SNP data. Genotype data can be read, filtered, and analysed within a single environment, without the need for prior data cleaning or setup with other software. *tidypopgen*’s *gen_tibble* object structure makes analysis efficient and intuitive, while standardised tidy grammar makes data manipulation clear.

Functionality within *tidypopgen* supports cleaning and merging datasets, basic descriptive statistics, multivariate analysis, clustering algorithms, and F-statistics, as well as integrating with existing tools for population genetic analyses in R. We use the Human Genome Diversity Project SNP dataset (Li et al., 2008) to show that a basic population genetic workflow can be executed in under 25 lines of code in a single environment using one file set, without the need to write superfluous outputs or change directories. By supporting data assembly through to data analysis, *tidypopgen* significantly streamlines workflows without compromising speed or functionality.

## Introduction

In the 21^st^ century the field of population genomics experienced a dramatic proliferation of data. As the cost of sequencing has reduced and computational power has increased, research has begun to leverage whole genome sequences and expansive genomic databases, such as the 1000 genomes project (1000 Genomes Project Consortium, 2015) and UK Biobank data (Sudlow et al., 2015). Vast amounts of genome-wide single-nucleotide polymorphism (SNP) data are now available for many organisms and are employed to address hypotheses across fields of evolutionary biology, conservation, anthropology, and beyond. Available tools for population genetic analysis have proliferated as the number and size of datasets has increased and so has the complexity of available data.

Plenty of existing software enables the analysis of such datasets, but pipelines often employ a combination of BASH (Free Software Foundation, 2022), python (Python Software Foundation, 2023), and R (R Core Team, 2024) to clean and reformat data. Pipelines are encumbered by switching between coding languages and directories, which can be time consuming and error prone. Previous solutions have provided workflow scripts for standard analyses (Moreno-Mayar, 2022), aiming to minimise time spent on data manipulation between different formats. Toolkits such as *FrAnTK* (Moreno-Mayar, 2022) can make data coercion more accessible, but illuminate a key problem: performing standard population genetic analyses requires at each stage multiple coding languages, additional libraries, numerous bespoke scripts, and many file outputs. A unified solution to this analytical tangle is lacking.

There are many advantages to the R environment for genetic analysis, which have been explored in depth elsewhere (Paradis et al., 2017), with the ability to utilise methods from different packages being its most alluring quality. Existing population genetics and phylogenetics packages within R (R Core Team, 2024), such as *adegenet* (Jombart, 2008; Jombart and Ahmed, 2011), *pegas* (Paradis, 2010), and *ape* (Paradis, Claude and Strimmer, 2004), have intuitive user interfaces with clear object structure and wide ranges of functionality. These methods allowed sufficient flexibility for subsequent packages, such as *dartR* (Gruber et al., 2018; Mijangos et al., 2022) and *SambaR* (de Jong et al., 2021), to build upon existing object structures and add functionality. Previous packages, such as *snpR* (Hemstrom and Jones, 2023), facilitate the incorporation of detailed multilevel metadata into genetic analysis by providing a data frame-like object structure, which also helpfully stores the results of statistical analyses on the given object.

However, existing methods can be slow, requiring users to run analyses on high performance clusters for sufficient speed (Gruber et al., 2018), and are sometimes memory intensive and lacking in scalability, placing limits upon the size of datasets that can be analysed. Packages may require the installation of many dependencies (de Jong et al., 2021), or conversion between object types to take advantage of their full functionality, which is both resource-consuming and potentially error-prone. Moreover, few packages offer functionality for basic data cleaning, meaning that most pipelines begin by using PLINK (Purcell et al., 2007) even when subsequent analysis is performed elsewhere.

We present *tidypopgen*, a new R package for population genetic analysis of SNP data which combines computational efficiency with ease of use. To streamline population genetics pipelines, *tidypopgen* creates a comprehensive approach entirely within the R environment. Data can be read into *tidypopgen* from vcf, packedancestry and PLINK formats, or can alternatively be constructed in R by providing a matrix of genotypes. The package then provides flexible ways to filter, analyse, and plot data. *tidypopgen* includes standard population genetic statistical analyses (e.g *F*_*ST*,_ *F*_*IS*_ and runs of homozygosity), options for multivariate analyses (multiple principal components analysis (PCA) methods, including projecting ancient DNA, as well as discriminant analysis of principal components), relatedness metrics (IBS and KING robust coefficients), and integration with the R package *Admixtools2* (Maier and Patterson, 2024) to calculate F-statistics.

### The *gen_tibble*

Large genetic datasets pose a challenge when coded as simple matrices, as they can be too large to store in memory. The packages *ape* (Paradis, Claude and Strimmer, 2004) and *adegenet* (Jombart and Ahmed, 2011), arguably the most popular for genetic analysis, employ data compression to avoid over-extending RAM, using two bits per diploid biallelic SNP to store 4 SNPs per byte. However, the ever-increasing size of genetic datasets demands new approaches to managing memory. In *tidypopgen*, we use the infrastructure developed by *bigstatsr* and *bigsnpr*, where data are stored in a file backed matrix (FBM) structure to keep large genotype files in a compressed matrix format on disk (Privé et al., 2018). Functions in *tidypopgen* use memory mapping, where algorithms do not load all the data in memory at once, but rather work sequentially on blocks of loci from the FBM and combine the intermediate results into a final set of quantities. This allows the analysis of very large datasets on computers with limited RAM. *tidypopgen* builds on this infrastructure further by creating the *gen_tibble* object. This is a subclass of tibble where the genotypes of individuals, loci information and metadata are stored as a special column that can be manipulated with familiar tidy grammar and clear verb-based commands, such as ‘*show_genotypes()*’ or ‘*select_loci()*’.

Following the tidyverse philosophy (Wickham et al., 2019), each row of a *gen_tibble* is an individual, with columns providing metadata. Each *gen_tibble* contains the compulsory columns ‘id’ (individual code) and ‘genotypes’. A link to the FBM representing the individuals’ genotypes is stored as an attribute of the ‘genotypes’ column of the *gen_tibble*. Loci information (that would be contained in a PLINK .bim or .map file) is also stored separately as an attribute. The use of the FBM infrastructure means that each *gen_tibble* has a set of backing files; a .gt file, containing all the metadata stored in the *gen_tibble*, together with an .rds and .bk file, containing the underlying genotype information. The *gt_save()* function saves *gen_tibble* objects at the end of each R session, ready to be reloaded with *gt_load()* at the beginning of the next session. It is possible to generate multiple *gen_tibbles* that use the same backing file set, where each *gen_tibble* represents a different subset of individuals or loci. This makes creating multiple versions or subsets of the same dataset easy and memory efficient: only one backing file set is required to save multiple subsets of the same data. For example, the *gen_tibble* objects saved as ‘example.gt’ and ‘example_ld_pruned.gt’ can use the same ‘example.rds’ and ‘example.bk’ backing files.

The *gen_tibble* structure therefore combines the best elements of efficient data storage, using *bigsnpr’s* FBM (Privé et al., 2018), with the flexibility of tibbles. The FBM structure also provides the capacity to store imputed genotypes efficiently, making it possible to create a single pipeline able to switch between imputed and non-imputed data, without writing any intermediate outputs. This means that, instead of imputing missing genotypes separately at each stage of analysis, data are imputed once only, ensuring replicability. The tibble structure also provides the opportunity to store plenty of per-individual metadata in each *gen_tibble* simply by appending columns. Users can group and ungroup a *gen_tibble*, as with a standard tibble, to run analyses that compare populations, or to allow flexibility in plotting data. This is analogous to the use of facets in *snpR* (Hemstrom and Jones, 2023) or ‘population’ arguments in *pegas* (Paradis, 2010). Users can also subset the *gen_tibble*, either to remove individuals for quality control or to run analyses on a specific sub-group of data.

Data can be exported from *tidypopgen* back into PLINK or .vcf format, allowing the user to switch between programs if necessary and making it easy to incorporate into existing workflows. *tidypopgen* can also convert a *gen_tibble* to the *genlight* and *genind* objects for *adegenet* (Jombart, 2008), or, additionally, to *heirfstat* objects (Goudet, 2005). This enables the use of functions already available within these packages, and within others based on these object structures (Gruber et al., 2018; de Jong et al., 2021; Mijangos et al., 2022).

### Workflow and examples

Documentation of functions and example workflow vignettes are available on the website (https://evolecolgroup.github.io/tidypopgen/index.html). *tidypopgen* has been tested with continuous integration across Windows, Mac, and Linux, and depends on R version 3.0.2 or higher (R Core Team, 2024).

The following examples use the Human Genome Diversity Project SNP dataset (Li et al., 2008), Malagasy data from Pierron et al. (2014), and modern and ancient data from European individuals taken from Lazaridis et al. (2016).

### Quality control

Basic data cleaning in *tidypopgen* is performed by two functions: *qc_report_loci()* and *qc_report_indiv(). qc_report_loci()* supplies a per-locus summary of the dataset, including rate of missingness, minor allele frequency, and a Hardy-Weinberg exact p-value for each SNP. These statistics can also be easily computed within each population. For example, if a *gen_tibble* contains multiple populations, Hardy-Weinberg p-values can be computed for each locus within each population by grouping the *gen_tibble* with *group_by()*, before running *qc_report_loci(). qc_report_indiv()* provides rate of missingness and observed heterozygosity for each individual. These outputs can be quickly visualised using built in *autoplot()* methods, exemplified in Fig. 2, inspired by the package *plinkQC* (Meyer, 2020).

**Figure 1:**
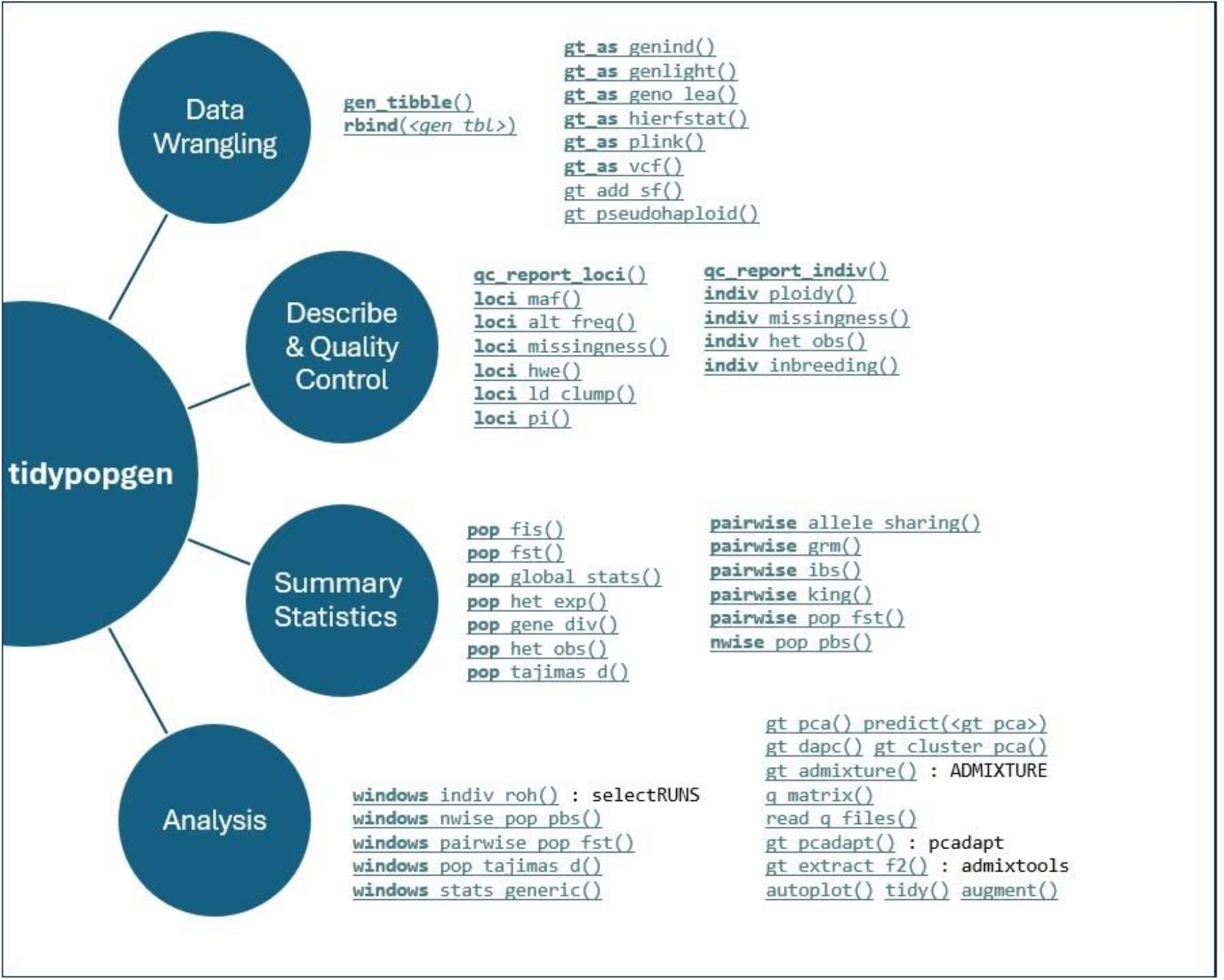
A schematic of key functions in the tidypopgen package. Functions that integrate with external packages are annotated with the package name.

**Figure 2:**
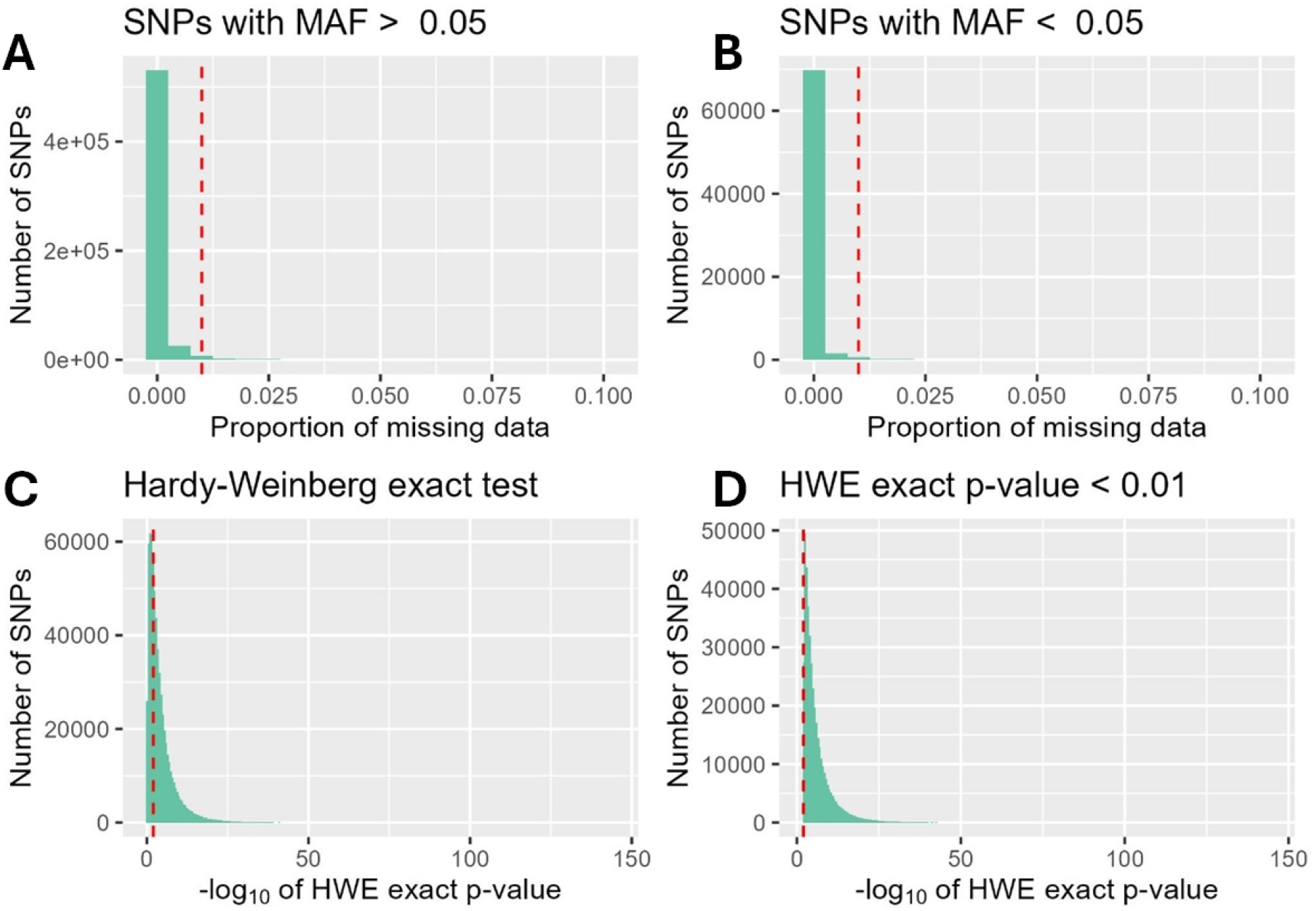
Quality control autoplots using HGDP data (<640000 SNPs). A) Histogram of the proportion of missing data for SNPs with minor allele frequency (MAF) over 0.05. B) Histogram of the proportion of missing data for SNPs with MAF under 0.05. C). Histogram of Hardy-Weinberg exact test (HWE) p-values. D) Histogram of significant HWE p-values.

The use of tidy grammar in *tidypopgen* means that data are filtered using the functions *select_loci()* and *filter()*, as if sub-setting a simple tibble. Example code for filtering data is given at https://evolecolgroup.github.io/tidypopgen/articles/a02_qc.html *tidypopgen* does not assume number of chromosomes and is therefore designed to handle biallelic autosomal SNP data from any organism. Sex chromosomes of the study organism can be easily removed after loading data using *select_loci_if()*. In addition to the standard filtering provided in the *qc_report* functions, related individuals can be identified using the *pairwise_king()* function, which calculates the KING kinship coefficient matrix (Manichaikul et al., 2010). Alternatively, the user can provide a ‘kings_threshold’ argument to *qc_report_indiv()*, to obtain an unrelated set based on the threshold provided. *tidypopgen* also uses the clumping method introduced by *bigsnpr* (Privé et al., 2018) to quality control for linkage disequilibrium with *loci_ld_clump()*.

### Merging

A key advantage of *tidypopgen* is its transparency of behaviour when merging data from multiple SNP arrays; a common challenge, particularly in human population genetics. Merging data requires multiple steps: mapping data to the same human genome build, resolving stranding inconsistencies, and removing any triallelic or ambiguous SNPs. In *tidypopgen, gen_tibble* objects are merged using the *rbind()* function, which automatically resolves strand inconsistencies and identifies ambiguous SNPs in a single command. Data can be merged either by rsID or by chromosome and position. Before merging data, the function *rbind_dry_run()* reports the overlap of the two datasets, the number of ambiguous SNPs, and, if requested, which SNPs require ‘flipping’ to handle strand inconsistencies. For example, when merging the HGDP data with Malagasy data from Pierron et al., (2014), *rbind_dry_run()* shows that the datasets overlap by 349,759 SNPs, with no ambiguous SNPs in either set, and that 63,927 SNPs in the latter data are flipped to match the reference HGDP strand. A full merge can then be performed easily with *rbind()*, generating a new *gen_tibble* object containing the merged dataset for further analyses.

### Imputation

*tidypopgen* implements the methods of *bigsnpr* (Privé et al., 2018) to provide fast imputation of missing genotypes using the functions *gt_impute_simple()*, with options to impute by mode, mean, or at random, and *gt_impute_xgboost()*, which uses a local XGBoost model. These approaches are suitable for small levels of missing data but should not be used for more substantial imputation (e.g. for low coverage aDNA data). The user can set and track whether a *gen_tibble* is using imputed data with the functions *gt_set_imputed(), gt_has_imputed()*, and *gt_uses_imputed()*. Functions that do not handle missingness will automatically use imputed data, whereas functions that manage missingness methodologically will not use the imputed dataset by default. Calculation of summary statistics, for example *F*_*ST*_ using *pairwise_pop_fst()*, will therefore only use the original dataset, unless the user explicitly requests to use the imputed data. The user can therefore control where imputed data is used throughout the pipeline and determine the method used for imputation.

### Summary statistics

Standard statistics for population genetic analyses are available in *tidypopgen*, with their grammar organised according to tidyverse principles. *loci_* functions operate over loci (e.g *loci_alt_freq()*) and *indiv_* functions operate over individuals (e.g *indiv_missingness()*). *pop_* functions (e.g *pop_fst()*) operate on a *gen_tibble* grouped by population, *pairwise_* functions (e.g *pairwise_ibs()*) compute pairwise statistics between pairs of individuals or pairs of populations, while *nwise_* functions (e.g *nwise_pop_pbs()*) compute statistics between all combinations of n individuals or populations. Furthermore, *windows_* functions (e.g *windows_pop_tajimas_d()*) calculate a per-locus statistic and compute a summary for each window, with arguments for users to define parameters (e.g. window size, overlap).

Ancient DNA datasets that contain pseudohaploid samples are handled by *gt_pseudohaploid()*, which assigns a specific genotype code to the *gen_tibble* and modifies the ploidy recorded for individuals. After assigning the genotype code, *tidypopgen* functions recognise that the *gen_tibble* contains pseudohaploid data and adapt algorithms accordingly. For example, the function *loci_alt_freq()* adjusts allele counts for pseudohaploid individuals, where dosages in the genotype matrix will be coded as either 0 or 2, but these denote only a single copy of the reference or alternate allele.

### Visualising

*tidypopgen* includes three methods for PCA; *gt_pca_autoSVD(), gt_pca_partialSVD()*, and *gt_pca_randomSVD(). gt_pca_autoSVD()* automatically implements initial LD clumping and removal of long-range regions of linkage, before running the PCA algorithm. The latter options allow the user to adjust the algorithm used based on dataset size, with guidance provided in the *gt_pca* documentation. Eigenvalues, loadings, and PCA scores are then stored in a *gt_pca* object, ready to *autoplot()* or tidy into a tibble structure for bespoke plotting. Fig. 3 visualises the merged HGDP and Pierron et al. (2014) data in a PCA, illustrating how the Malagasy samples form a cline between African and Asian populations across the first principal component.

**Figure 3:**
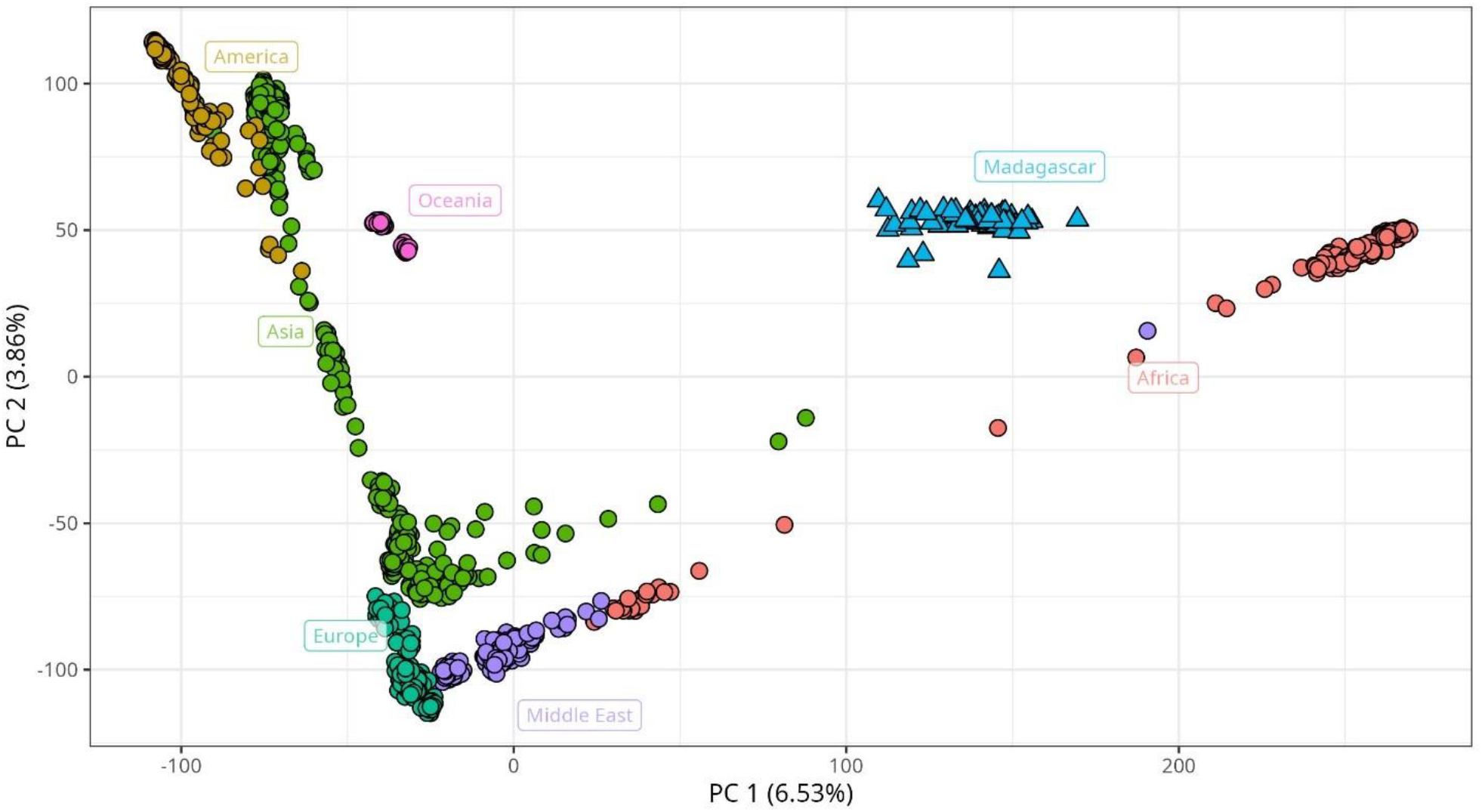
Principal components analysis of HGDP and Malagasy data taken from Pierron et al. 2014. The 69 Malagasy individuals cluster together between African and Asian populations across the first principal component.

New data can be projected onto an existing *gt_pca* object using the *predict()* function, where the PC scores of new samples are calculated from the precomputed PCA. This function offers the methods ‘least_squares’ (as implemented by *SMARTPCA* (Patterson, Price and Reich, 2006)), ‘simple’ and ‘OADP’ (Online Augmentation, Decomposition, and Procrustes projection), (both described in Zhang, Dey and Lee (2020)). Fig. 4 uses data from Lazaridis et al. (2016) to demonstrate this capacity; ancient samples are projected onto a PCA of modern west Eurasian individuals using the ‘least_squares’ projection method. *gt_pca* objects can also be used to perform Discriminant Analysis of Principal Components (DAPC) (Jombart, Devillard and Balloux, 2010) with *gt_dapc()*, for characterising population structure through clustering. The function *gt_cluster_pca_best_k()* then provides options for identifying the optimum clustering level.

**Figure 4:**
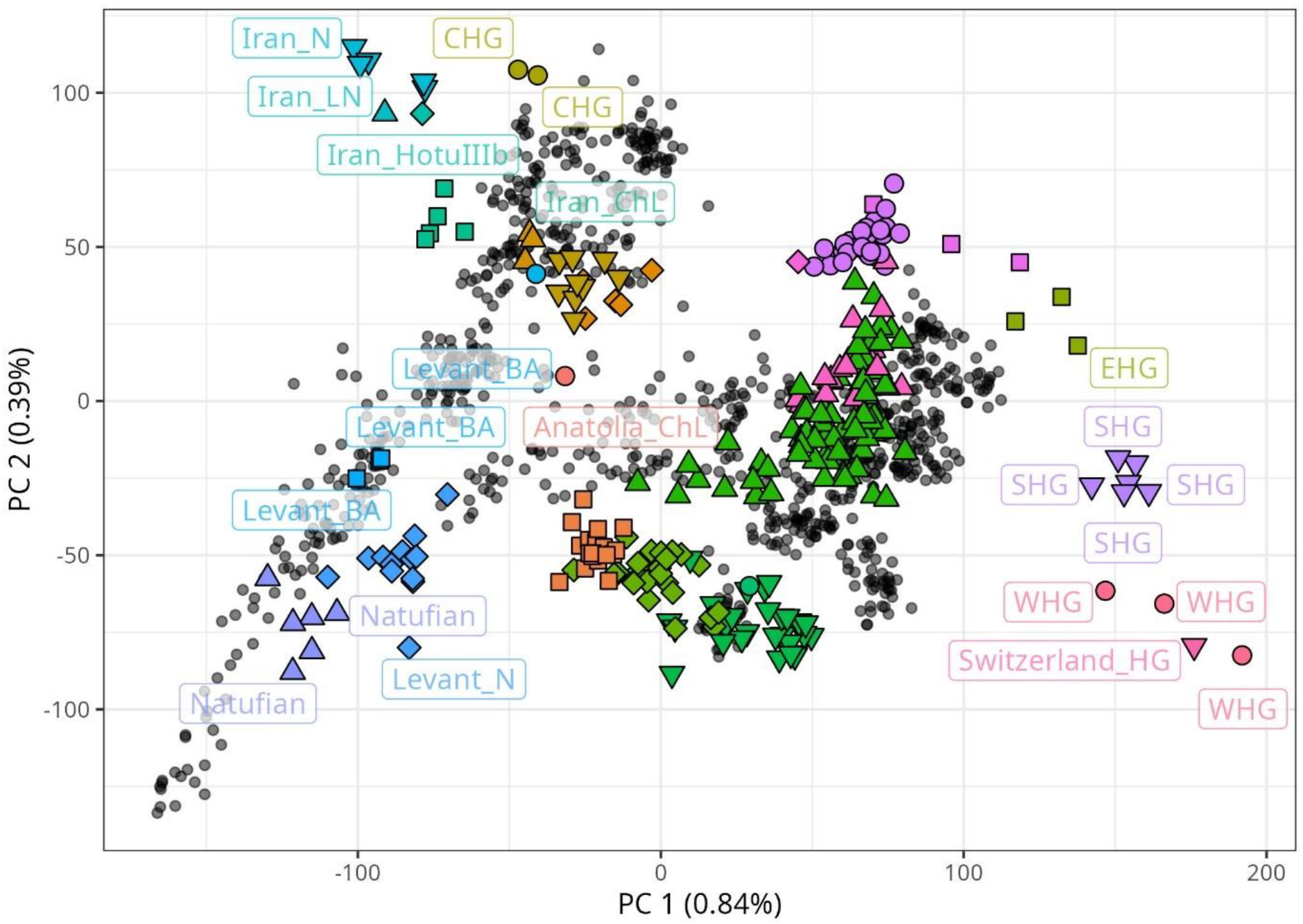
Principal components analysis using data from Lazaridis et al. (2016). Principal components are calculated using 991 modern west Eurasian individuals (grey points). 278 Ancient individuals (larger coloured points) are projected.

For further clustering analysis, *tidypopgen* relies on external software, but provides functions to prepare data and run clustering in the background. Currently, ADMIXTURE (Alexander, Novembre and Lange, 2009) is run through *gt_admixture()*, while *gt_snmf()* exports data to *LEA* (Frichot and François, 2015) for sparse nonnegative matrix factorization. Both options return *gt_admix* objects, which can be visualised through *autoplot()*. Additionally, .Q matrix files from any alternative workflow run outside of R can be easily read into *tidypopgen*, tidied, and plotted using *read_q_files()*.

*tidypopgen* integrates with the *Admixtools2* R package for easy calculation of F-statistics (Maier and Patterson, 2024). After reading and filtering data, the function *gt_extract_f2()* computes allele frequencies and *F*_*2*_ statistics for population pairs, using the same method as *Admixtools2*, directly from a *gen_tibble. Admixtools2* functions can then be directed to the precomputed *F2* folder and used to calculate more complex F-statistics.

### Benchmarking

To exemplify the performance of *tidypopgen*, we ran a basic analysis using HGDP SNP data (1,043 individuals and ∼570,000 SNPs). The whole analysis requires 24 lines of code: from data upload through cleaning, PCA, DAPC, pairwise *F*_*ST*_ across 50 populations, to writing output files. It takes less than 38 seconds on a laptop and less than 18 seconds on a powerful desktop. Replication in *dartR* (Gruber et al., 2018; Mijangos et al., 2022) and *snpR* (Hemstrom and Jones, 2023) was not possible, as the size of the dataset caused memory allocation errors. Full details are available on the *tidypopgen* website (https://evolecolgroup.github.io/tidypopgen/articles/benchmark_hgdp.html).

## Conclusion

*tidypopgen* integrates all stages of population genetic analysis into a cohesive framework, using the R environment. Large datasets are managed efficiently with the FBM-backed *gen_tibble* object structure, making standard analyses swift even without high-performance computing equipment. By removing the common time sinks of downloading and learning multiple suites of software, merging datasets, and wrangling outputs, *tidypopgen* streamlines workflows and frees time for analysis.

## Author contributions

A.M, and E.C designed the package functionality with input from E.T and J.H. A.M and E.C wrote the R package and documentation. A.M, E.T, and E.C wrote unit tests. E.C led the writing of the manuscript with input from A.M, J.H, and E.T.

## Acknowledgements

We thank Margherita Colucci, Aramish Fatima, Anahit Hovhannisyan, Cecilia Padilla-Iglesias and Andrea Pozzi for testing the package while under development and providing feedback. We thank Michela Leonardi for comments on the manuscript, and Max Brown for discussion on vcf parsing.

## Data availability

HGDP data used in Fig 1. and Fig. 3 is available on Zenodo at https://doi.org/10.5281/zenodo.15582364. Ancient and modern data taken from Lazaridis et al. (2016). used in Fig. 4 are available on the Reich lab website under Datasets https://reich.hms.harvard.edu/datasets The Malagasy data taken from Pierron et al. (2014) used in Fig. 3 are available online in the Gene Expression Omnibus (GEO) database, using the accession number GSE53445.

